# Demography and Selection Shape Transcriptomic Divergence in Field Crickets

**DOI:** 10.1101/193839

**Authors:** Thomas Blankers, Sibelle T. Vilaça, Isabelle Waurick, David A. Gray, R. Matthias Hennig, Camila J. Mazzoni, Frieder Mayer, Emma L. Berdan

**Affiliations:** Behavioural Physiology, Department of Biology, Humboldt-Universität zu Berlin, Berlin, Germany, D-10115; Museum für Naturkunde Berlin, Leibniz Institute for Evolution and Biodiversity Science, Berlin, Germany, D-10115; Current Address: Department of Neurobiology and Behavior, Cornell University, Ithaca, NY, USA, 14853; Berlin Center for Genomics in Biodiversity Research (BeGenDiv), Berlin, Germany, D-14195; Leibniz-Institut für Zoo- und Wildtierforschung (IZW), Berlin, Germany, D-10315; Department of Biology, California State University Northridge, Northridge, CA, USA, 91330-8303; Berlin-Brandenburg Institute of Advanced Biodiversity Research (BBIB), Berlin, Germany, D-14195; Current Address: Department of Marine Sciences, University of Gothenburg, Gothenburg, Sweden, SE - 40530

**Keywords:** gene flow, mating behavior, *Gryllus*

## Abstract

Gene flow, demography, and selection can result in similar patterns of genomic variation and disentangling their effects is key to understanding speciation. Here, we assess transcriptomic variation to unravel the evolutionary history of *Gryllus rubens* and *Gryllus texensis*, cryptic field cricket species with highly divergent mating behavior. We infer their demographic history and screen their transcriptomes for footprints of selection in the context of the inferred demography. We find strong support for a long history of bidirectional gene flow, which ceased during the late Pleistocene, and a bottleneck in *G. rubens* consistent with a peripatric origin of this species. Importantly, the demographic history has likely strongly shaped patterns of neutral genetic differentiation (empirical *F_ST_* distribution). Concordantly, *F_ST_* based selection detection uncovers a large number of outliers, likely comprising many false positives, echoing recent theoretical insights. Alternative genetic signatures of positive selection, informed by the demographic history of the sibling species, highlighted a smaller set of loci; many of these are candidates for controlling variation in mating behavior. Our results underscore the importance of demography in shaping overall patterns of genetic divergence and highlight that examining both demography and selection facilitates a more complete understanding of genetic divergence during speciation.

## INTRODUCTION

The study of speciation and the origins of earth’s biodiversity are at the core of evolutionary biology. An important first step is understanding the mechanisms that drive genetic divergence between closely related groups of organisms. In the age of next-generation sequencing, our understanding of these mechanisms is rapidly advancing. However, a variety of processes such as gene flow, local variation in recombination and mutation rates, linked or background selection, and divergent selection often simultaneously influence genetic variation between diverging lineages and the different processes may leave similar signatures in the genome (Noor and Bennett 2009; Feder et al. 2012; Nachman and Payseur 2012; Cutter and Payseur 2013; Seehausen et al. 2014; Burri et al. 2015). Therefore, to understand how populations diverge, how reproductive isolation evolves, and how this affects the genome, it is essential that we examine both selective and neutral processes.

Recently, the role of gene flow in speciation has drawn renewed attention (Smadja and Butlin 2011; Feder et al. 2013; Sousa and Hey 2013; Servedio 2015; Ravinet et al. 2017). It was once thought that reproductive barriers could only evolve in allopatry (Mayr 1963; Bolnick and Fitzpatrick 2007). However, this view has shifted due to accumulating evidence for varying rates of gene flow during early divergence (Bolnick and Fitzpatrick 2007; Nosil 2008; Bird et al. 2012). Although ‘true’ sympatric speciation is likely rare, there is nowadays a general acceptance that some amount of gene flow occurs during many speciation events, i.e. parapatric speciation (Coyne and Orr 2004; Smadja and Butlin 2011; Arnold 2015).

Speciation with gene flow has attracted special attention because strong divergent selection in combination with high migration rates may lead to higher (than background) genomic divergence in the regions harboring loci important for reproductive isolation and local adaptation (Turner et al. 2005; Nosil et al. 2009; Cutter and Payseur 2013; Feder et al. 2013; Ravinet et al. 2017). However, variation in levels of divergence across the genome may also strongly depend on locally reduced intraspecific diversity due to demographic effects or variation in mutation and recombination rates (Nachman and Payseur 2012; Cruickshank and Hahn 2014; Burri et al. 2015). Additionally, the likelihood of detecting the effects of selection above background levels of genomic variation is highly dependent on the genetic architecture of the traits under selection (Jiggins and Martin 2017) and the strength of selection (Ortiz-Barrientos and James 2017). These caveats warrant caution in the interpretation of the results from genomic scans, especially without a detailed understanding of the behavioral ecology and evolutionary history of the study system (Ravinet et al. 2017). Thus, a primary goal of studies aiming to elucidate the effects of selection on genetic variation should be to consider patterns left by neutral or demographic processes that could occlude the genomic signature of selection.

Here, we bring this goal into practice by inferring demographic history and characterizing the resulting patterns of genetic variation in the absence of selection. We then use the resulting neutral expectation to inform our inference of putative signatures of selection. For this approach, we use transcriptomic data from two sexually isolated field cricket species, *Gryllus rubens* and *Gryllus texensis*. Given their current distributions (Fig. 1), it is likely that interspecific gene flow has played a dominant role in the evolutionary history of *G. texensis* and *G. rubens*; although, contemporary gene flow is unlikely based on lack of genetic or phenotypic evidence supporting hybridization in nature (Walker 1998; Gray and Cade 2000; Higgins and Waugaman 2004) and reinforcement (Gray and Cade 2000; Izzo and Gray 2004). Additionally, a mitochondrial study found evidence that suggests *G. rubens* has a peripatric origin from *G. texensis* (Gray et al. 2008) and thus divergence between *G. texensis* and *G. rubens* may be associated with a strong bottleneck for the latter but not the former species.

**Fig. 1.**
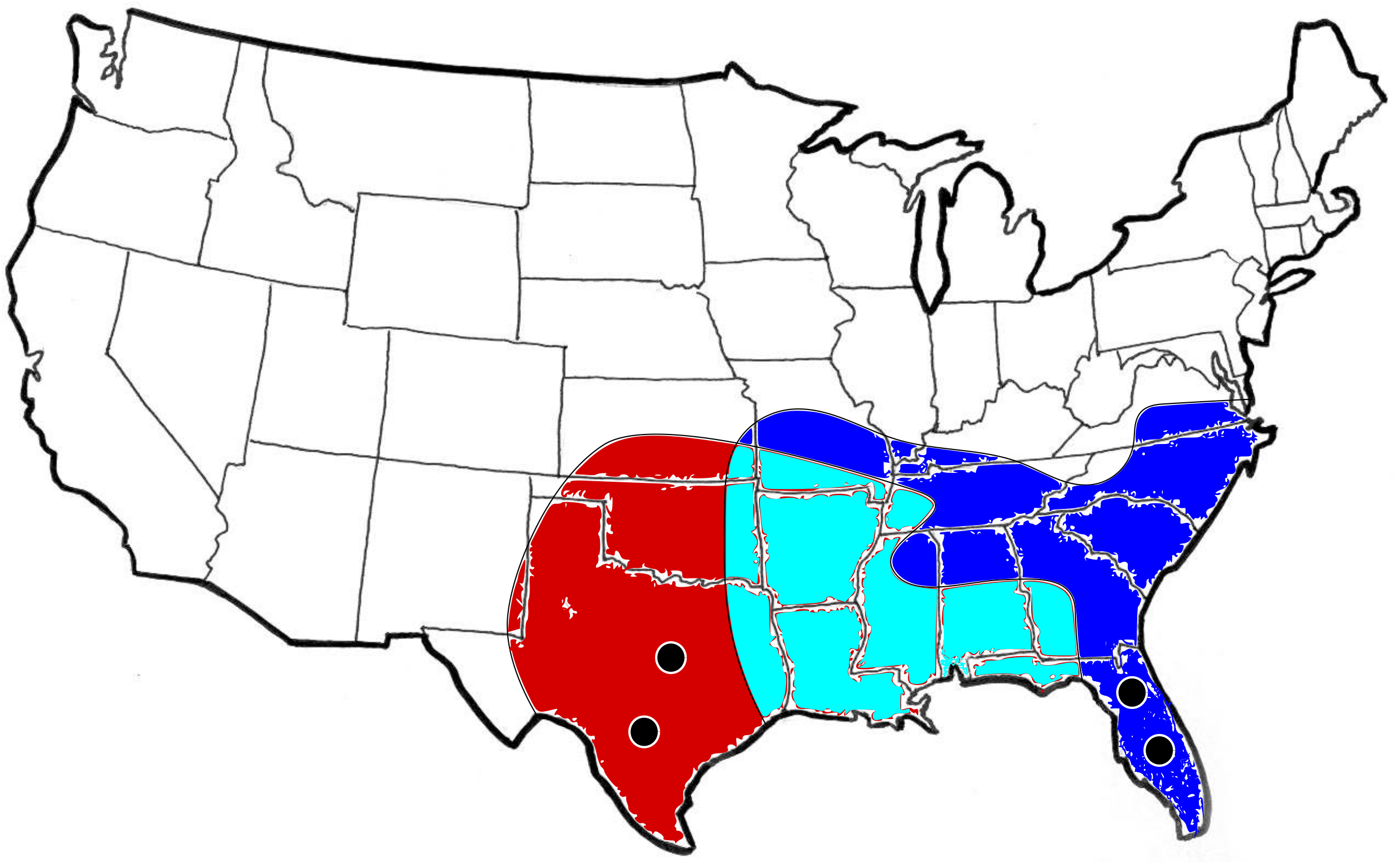
Geographic distributions for *G. texensis* (red) and *G. rubens* (blue). The sympatric zone is marked with turquoise. The distributions are approximate and based on the Singing Insects of North America data base (http://entnemdept.ufl.edu/Walker/buzz/). The black dots in Texas and Florida represent the sampling locations for *G. texensis* and *G. rubens*, respectively.

In addition to demographic processes selection also likely played a role in the divergence of *G. texensis* and *G. rubens*. There is striking variation in acoustic sexual communication behavior in this system, involving multiple traits that compose the wing-generated calling song produced by cricket males and corresponding female preferences. This implies a strong selective pressure on genes related to mating signals. Variation in the cricket mating song depends on (i) the morphology and resonant properties of the wings, (ii) neural networks called central pattern generators that control rhythmic wing movement, and (iii) neuromuscular properties of the muscles that affect the temporal rhythm of the song (reviewed in Gerhardt and Huber 2002). Similarly, song recognition and preference in females are controlled by a complex network of neurons and likely depend on properties of ion channels, in particular potassium channels mediating inhibitory effects (Hennig et al. 2014; Schoneich et al. 2015; Göpfert and Hennig 2016). If there is indeed a strong selective pressure on mating behavior in this system, selection signatures are expected to be biased towards gene products that affect the properties of muscles, neuromuscular junctions, and neurotransmitter activity related to rhythmic behaviors and perception, as well as mating behavior variation more broadly. Importantly, combining the inference of putative selection signatures with the demographic analyses allows us to interpret the perceived effects from selection in the appropriate historical context and make predictions about the joint effects from neutral and selective forces during population divergence.

## MATERIALS & METHODS

### Study system

*Gryllus texensis* and *G. rubens* are widely distributed across the southern Gulf and Mid-Atlantic States in North America, with a broad sympatric region from eastern Texas through western Florida (Fig. 1). Males are morphologically cryptic (Gray et al. 2008) and there is no documented ecological divergence (Gray 2011). However, females differ in the length of the ovipositor (Gray et al. 2001), which tentatively reflects ecological adaptation to different soil types (Bradford et al. 1993). In nature, divergence in acoustic signals and preferences is a strong premating barrier acting through both species-specific long-distance mate attraction songs (Walker 1998; Gray and Cade 2000; Blankers et al. 2015a) and close-range courtship songs (Gray 2005; Izzo and Gray 2011). Reproductive isolation is maintained in the zone of overlap, but there is no evidence for reproductive character displacement, indicating that reinforcement is unlikely to affect divergence in these species (Higgins and Waugaman 2004; Izzo and Gray 2004).

### Sample collection

Animals were collected in the USA in Lancaster and Austin (TX; ca. 80 *G. texensis* females) and in Lake City and Ocala (FL; ca. 40 *G. rubens* females) in autumn 2013 (Fig. 1 black dots). Collected females, which are typically already inseminated in the field, were housed in containers in groups of up to 15 individuals with gravel substrate, shelter, and water and food *ad libitum*. Each container also contained a cup with vermiculite for oviposition. During two weeks, eggs were collected and transferred to new containers; hatchlings were then reared to adulthood. We used laboratory-raised offspring of the field-caught females between one and three weeks after their final molt rather than field-caught specimens to standardize rearing conditions across all samples. All animals (males and females) were played back an artificial stimulus resembling the conspecific male song for 10 minutes prior to sacrificing the animal. The rationale here was that one of our primary objectives was to look at genetic divergence in relation to mating behavior polymorphism. In case specific genes involved in female preference behavior were only expressed upon hearing a male song signal, this could potentially be overcome by a brief play back 30 – 120 minutes prior to RNA preservation. Stimulus play back occurred for females and males to standardize the RNA sampling method across sexes. Within two hours of stimulus presentation, we sacrificed the cricket, removed the gut and then preserved the body in RNAlater following the manufacturer’s instructions; samples were then stored at −80 °C until RNA isolation. A total of five males and five females were used from each of the two populations for each species (40 individuals in total; randomly sampled across containers when there were multiple containers for crickets from the same population). Total RNA extraction and directional, strand-specific Illumina library preparation were done as described in a recently published transcriptomic resource for *Gryllus rubens* (Berdan et al. 2016).

### SNP calling

Raw reads were processed using Flexbar (Dodt et al. 2012) to remove sequencing primers, adapters, and low quality bases on the 3’ end of the individually barcoded reads. Samples were mapped to the *G. rubens* reference transcriptome (Berdan et al. 2016) using Bowtie2 (Langmead and Salzberg 2012) with default parameters but specifying read groups to mark reads as belonging to a specific individual. Duplicate reads were marked using ‘picard’ (http://broadinstitute.github.io/picard). The Genome Analysis Toolkit (GATK, DePristo *et al*. 2011; Van der Auwera *et al*. 2013) was used to call genotypes with the GATK-module ‘UnifiedGenotyper’(Van der Auwera et al. 2013). The variants were then filtered to only retain high quality SNPs based on the recommendations on the GATK website (https://gatkforums.broadinstitute.org/gatk/discussion/comment/30641, accessed on 05/05/2015) and as described in a previous study (Berdan et al. 2015). The minor allele frequency (MAF) cut-off was set at 0.025 (a minimum of two copies of the allele).

Our sampling design was optimized to standardize the conditions under which we stored RNA samples, but potentially introduced a bias towards collecting related individuals. This may affect both demographic inference and the summary statistics used to identify selective sweeps. To correct for the potential cryptic relatedness, we used the PLINK methods-of-moments approach (Purcell et al. 2007) implemented in the SNPrelate package (Zheng et al. 2012) in R (R Development Core Team 2016) to estimate kinship coefficients for all pairs based on the allele frequencies within each population sample. We excluded eight individuals that showed estimated kinship coefficients above 0.125 (half-sib level) with other individuals from their population, leaving 17 *G. texensis* and 15 *G. rubens* individuals for all downstream analyses.

### The demographic history

We first tested whether the sampled populations show geographic genetic structure. We inspected allele frequency variation within and between species and populations using principal component analysis. We also ran STRUCTURE (Falush et al. 2003), once for each species separately and once combining the species, using a single SNP locus per contig (8,835 randomly drawn SNPs). We used the admixture model with sampling location as prior information. We ran STRUCTURE with an MCMC chain length of 100,0 and with a burn-in length of 10,000 for K=1 through K=5 (K=4 for the species-specific runs) with three repetitions for each K-value. Results were analyzed using STRUCTURE HARVESTER (Earl and vonHoldt 2012) using the log-likelihood to compare K=1 versus all other values for K and the delta K method (Evanno et al. 2005) to compare K=2 versus all higher values of K.

To investigate the demographic history of *G. rubens* and *G. texensis*, we used the approximate Bayesian computation framework (ABC, Beaumont *et al*. 2002). We used ABCsampler from the ABCtoolbox package (Wegmann et al. 2009) to simulate our data under different demographic scenarios in fastsimcoal v2.5.2.3 (Excoffier and Foll 2011; Excoffier et al. 2013) and to calculate summary statistics using arlsumstat v.3.5.1.3 in Arlequin v 3.5 (Excoffier and Lischer 2010). We performed the analysis using the sequences from 1000 randomly drawn contigs (not including contigs with zero SNPs), using fixed recombination and mutation rates (both 1e-8) and the same minor allele frequency cut-off for the simulated data as for the observed data (0.025). To ensure that a MAF of 0.025 for the observed data was still maintained after removing the potentially related individuals, we removed all singletons that arose *post* subsampling. We initially calculated all between population summary statistics supported by arlsumstat. Then, using partial least squares regression (PLS), we retained the summary statistics with the highest predictive power (*i.e*. those with high factor loadings on the PLS components that significantly increase the predictive power of parameter estimates) for demographic estimates: the between-species mean and standard deviation of the number of polymorphic sites, the number of private polymorphic sites, Tajima’s D, and nucleotide diversity (π) in each species, as well as pairwise (between species) *F_ST_* and π. All statistics were calculated as averages across contigs.

We compared four possible (groups of) models: a simple divergence model (DIV; 4 parameters for population sizes and the timing/magnitude of demographic events), three models involving gene flow (either continuous, ancient or recent gene flow/secondary contact; CGF, AGF, RGF; 6-7 parameters), three models involving a bottleneck (for either or both species; RB, TB, BB; 6-8 parameters), and a model combining the most likely gene flow and most likely bottleneck model (AGFRB; 9 parameters). We intentionally considered only relatively simple models with few parameters to avoid the risk of overparameterization (Csilléry et al. 2010). For each model, we ran 200,000 iterations to do model selection. Prior ranges for population sizes and time points were chosen on a log-uniform scale spanning across several orders of magnitude and for bottleneck size and migration rates on a uniform scale not overlapping zero (Table 1).

**Table 1.** ABC estimates. Prior distributions (log-scale), posterior predictive checks and posterior parameter estimates (log scale, median and 95% highest posterior density interval) for the model are shown.

| Parameter | Prior<sup>a</sup> |  | Validation |  | Posterior |  |  |
| --- | --- | --- | --- | --- | --- | --- | --- |
|  | minimum | maximum | R<sup>2</sup> | RMSEP | 2.5% | Median | 97.5% |
| LOG<sub>10</sub>(N<sub>ANC</sub>) | 4.0 | 6.0 (lu) | 0.13 | 0.93 | 4.68 | 5.34 | 5.99 |
| LOG<sub>10</sub>(N<sub>RUB</sub>) | 3.0 | 6.0 (lu) | 0.90 | 0.32 | 4.03 | 4.50 | 4.70 |
| LOG<sub>10</sub>(N<sub>TEX</sub>) | 3.0 | 6.0 (lu) | 0.75 | 0.50 | 4.51 | 4.78 | 4.87 |
| LOG<sub>10</sub>(T<sub>SPLIT</sub>)<sup>b</sup> | 5.0 | 7.0 (lu) | 0.02 | 0.99 | 4.86 | 6.19 | 7.16 |
| LOG<sub>10</sub>(T<sub>ISO</sub>)<sup>b</sup> | 3.0 | 7.0 (lu) | 0.90 | 0.32 | 4.20 | 4.55 | 4.76 |
| LOG<sub>10</sub>(T<sub>BOT</sub>)<sup>b</sup> | 5.0 | 7.0 (lu) | 0.48 | 0.72 | 4.42 | 5.01 | 6.16 |
| BOTSIZE | 0.01 | 0.5 (u) | 0.16 | 0.91 | -0.04 | 0.15 | 0.48 |
| M<sub>TEX>>RUB</sub> | 0.01 | 0.5 (u) | 0.06 | 0.97 | 0.01 | 0.18 | 0.54 |
| M<sub>RUB>>TEX</sub> | 0.01 | 0.5 (u) | 0.06 | 0.97 | 0.01 | 0.27 | 0.74 |

<sup>a</sup> priors are uniformly (u) or log-uniformly (lu) distributed and do not overlap zero for migration rates and bottleneck size.
<sup>b</sup> the timing of demographic events is in (logarithm of) number of generation and both species have two generations annually.

After simulating the scenarios, model selection and posterior predictive checks were performed in R. Because of their similarity, the three bottleneck models and the three gene flow models were treated as two groups of models that were first tested inter-se; the best model of each group was then tested against the DIV and best combined models. We first retained the 1% of the samples that had the smallest Euclidean distance between the summary statistics of the simulated data and the observed data (‘ 1% nearest posterior samples’ from hereon) for each scenario separately. We then obtained a set of linear discriminants that maximized the distance among models within the nested categories (gene flow and presence of bottleneck). Next, posterior model probabilities were calculated based on these linear combinations of summary statistics using the ‘postpr’ function in the ‘abc’ package (Csilléry et al. 2012). The first two model selection steps were used to retain one gene flow and one bottleneck model with the highest posterior probability (‘best model’ from hereon). A third round of model selection was used to select among a simple divergence scenario (DIV), the best gene flow and bottleneck scenarios (AGF and RB, respectively; see Results), and a scenario combining the best gene flow and the best bottleneck scenario (AGFRB). Model selection was validated by performing leave-one-out cross validation with logistic regression using the ‘cv4postpr’ function. Here, one simulated sample, chosen at random from the posterior distribution, is left out and considered to be the “true” model while repeating the model selection step (with the remaining posterior samples) to evaluate the robustness of the model selection (Csilléry et al. 2012).

To estimate demographic parameters, we then ran 1,000,000 new simulations under the model(s) with the highest posterior probability. Posterior predictive checks were performed by calculating the predicted R^2^ and root mean squared error prediction (RMSEP) using the ‘pls’ package (Mevik and Wehrens 2007). We also used the ‘cv4abc’ function from the ‘abc’ package to evaluate prediction error. We estimated the demographic parameters with the ‘abc’ function using non-linear regression and a tolerance rate of 0.05.

An important goal of this study was to assess the effects of demography, in particular the timing of gene flow, on the *patterns* of transcriptome-wide genetic variation (*e.g*. the *F_ST_* distribution), rather than only on summary statistics. This will provide important insight into the extent to which loci that have evolved in the absence of selection are expected to confound the signatures of selection. We thus estimated a null distribution of the allele frequency spectrum (*i.e*. Tajima’s D, Tajima 1989) under the best fitting demographic model (see below). In addition, for the 1% nearest posterior samples of the models simulating continuous, recent, and ancestral gene flow and the AGFRB model we obtained the simulated *F_ST_* distribution for each posterior sample. The median and variation of these distributions were then visually contrasted with the observed *F_ST_* distribution.

### The role of selection

To assess the role of selection in driving genetic divergence, we employ three approaches that differ in their sensitivity to distinguish signals of selection from the confounding effects from past demographic events. All else being equal, variation in allele frequencies between populations is expected to increase more rapidly in the presence of selection. However, the most common measure of the variance in allele frequencies among populations, *F_ST_*, which is also a common test statistic to distinguish selected loci from the genomic background, has been criticized as a reliable indicator from various angles (*e.g*. Narum and Hess 2011; Cruickshank and Hahn 2014; Lotterhos and Whitlock 2014). Other methods may be better suited for detection of selected loci given strong demographic effects. For instance given sufficiently long divergence times and high levels primary or secondary gene flow, elevated sequence divergence (*d_xy_*) may be expected to better contrast the regions harboring loci involved in reproductive isolation from the rest of the genome (Nachman and Payseur 2012; Cruickshank and Hahn 2014). Additionally, a recent selective sweep may increase between population differentiation and decrease within population diversity and shift allele frequency spectrum (AFS) towards a higher frequency of rare alleles. Although demographic effects may also shift the AFS, these effects can be modeled and taken into account. Here, we contrast an *F_ST_* outlier scan (the “*F_ST_* approach” from hereon) with two alternative methods that should be better suited to withstand demographic effects (hereafter “d_xy_ approach” and the “selective sweep approach”, respectively).

We considered loci to be potentially under positive or divergent selection if they exceeded genomic background levels of (1) *F_ST_*, (2) absolute sequence divergence *(d_x_y)*, or (3) frequencies of rare alleles (Tajima’s D), low diversity (π), and high differentiation (*F_ST_*). For the *F_ST_* approach, we used the hierarchical island model (Slatkin and Voelm 1991) implemented in Arlequin (Excoffier et al. 2009; Excoffier and Lischer 2010). To accommodate the data to Arlequin’s input file restrictions, we only considered SNPs with MAF > 5% (81,125 SNPs). We pooled the two *G. rubens* populations in one group and two *G. texensis* populations in another group and performed 100,000 simulations to establish the neutral expectations for the relationship between among population heterozygosity and *F_ST_*. We considered all loci with *F_ST_* higher than the 99^th^ quantile for a given level of heterozygosity to be selection outliers.

For the d_xy_ approach and the selective sweep approach, we used VCFtools (Danecek et al. 2011) to calculate the following summary statistics: Tajima’s D (Tajima 1989), nucleotide diversity π (Nei and Li 1979), and weighted *F_ST_* (Weir and Cockerham 1984) in 1000 bp windows, and the absolute difference between the frequency of the major allele in the two species. We also calculated the average interspecific pairwise distance *d_xy_* for each window as *d_xy_* = π/(1-*F_ST_*), where π is the mean of the nucleotide diversity across species and *F_ST_* is the weighted mean *F_ST_* (Hudson et al. 1992; note that this method is similar to the often used *d_xy_* = *p_i_*(1-*p_j_*) + *p_j_*(1-*p_i_*), with *p_i_* and *p_j_* are the major or minor allele frequencies in species *i* and *j*, averaged across windows, weighed by the number of SNPs). For the d_xy_ approach, we retained the top 1% contigs with respect to *d_xy_* predicting that these loci have diverged relatively early in the evolutionary history and remained shielded from gene flow throughout. For the selective sweep approach, we retained all loci that had Tajima’s D below the 5% lowest simulated Tajima’s D values under the inferred demographic scenario and with values for π and *F_ST_* in the lowest and highest 10%, respectively. As this approach uses intraspecific population genetic data, we retained sets of outlier loci for both species separately

For all sets of outliers we checked for enriched Gene Ontology terms using ‘topGO’ (Alexa and Rahnenfuhrer 2016), part of the Bioconductor toolkit in R. The GO annotation was obtained from the *G. rubens* reference transcriptome (Berdan et al. 2016), which used the GO mapping module in Blast2Go (Conesa et al. 2005). We limited our gene set enrichment to biological process terms only and used the parent-child algorithm (Grossmann et al. 2007) to correct the *P* values for the ‘inheritance problem’ (i.e., the problem that higher GO terms inherit annotations from more specific descendant terms leading to false positives). We considered any GO term significantly enriched if the false discovery rate (Benjamini and Hochberg 1995) associated with the corrected P-value was below 10%. To get a more detailed picture of the putative functions of a given outlier locus, we looked up the functional annotation for the corresponding predicted gene product *(i.e*. the homolog with the highest similarity) on Flybase (Gramates et al. 2017) if that locus had been annotated using the *Drosophila melanogaster* proteome (see Berdan et al 2016 for details regarding transcriptome annotation).

## RESULTS

### Transcriptomic divergence

We sequenced RNA from 40 individuals (20 *G. rubens* and 20 *G. texensis)* on a HiSeq 2000 (Illumina, San Diego, CA, USA) obtaining on average 51,046,578 100-bp reads per individual (range 37,887,468-72,304,968) at a sequencing depth of eight libraries per lane. Reads mapped to the *G. rubens* transcriptome at an average rate of 83.2% (Table S1). Mapping rates were not higher in *G. rubens* despite the use of the *G. rubens* transcriptome *(G. rubens:* 83.0%; *G. texensis:* 83.0%; *P* = 0.9968), but females mapped at a significantly higher rate than males (86.0% versus 79.6%; *P* < 0.0001). At a MAF cut-off of 0.025 we found a total of 175,244 SNPs across 8835 contigs. The average transition-transversion ratio was 1.6:1. Nucleotide diversity (π) was similar among *G. rubens* (π = 0.11, σπ = 0.14) and *G. texensis* (π = 0.13, σπ = 0.15). Median *D* was 0.07 (first quantile: 0.05, third quantile 0.20) and 2.7% of the SNPs (4,828) were fixed between the species (Fig. 2A). Average Tajima’s D was negative for both species, but the distribution across loci showed substantial variation (Fig. 2B, C).

**Fig. 2.**
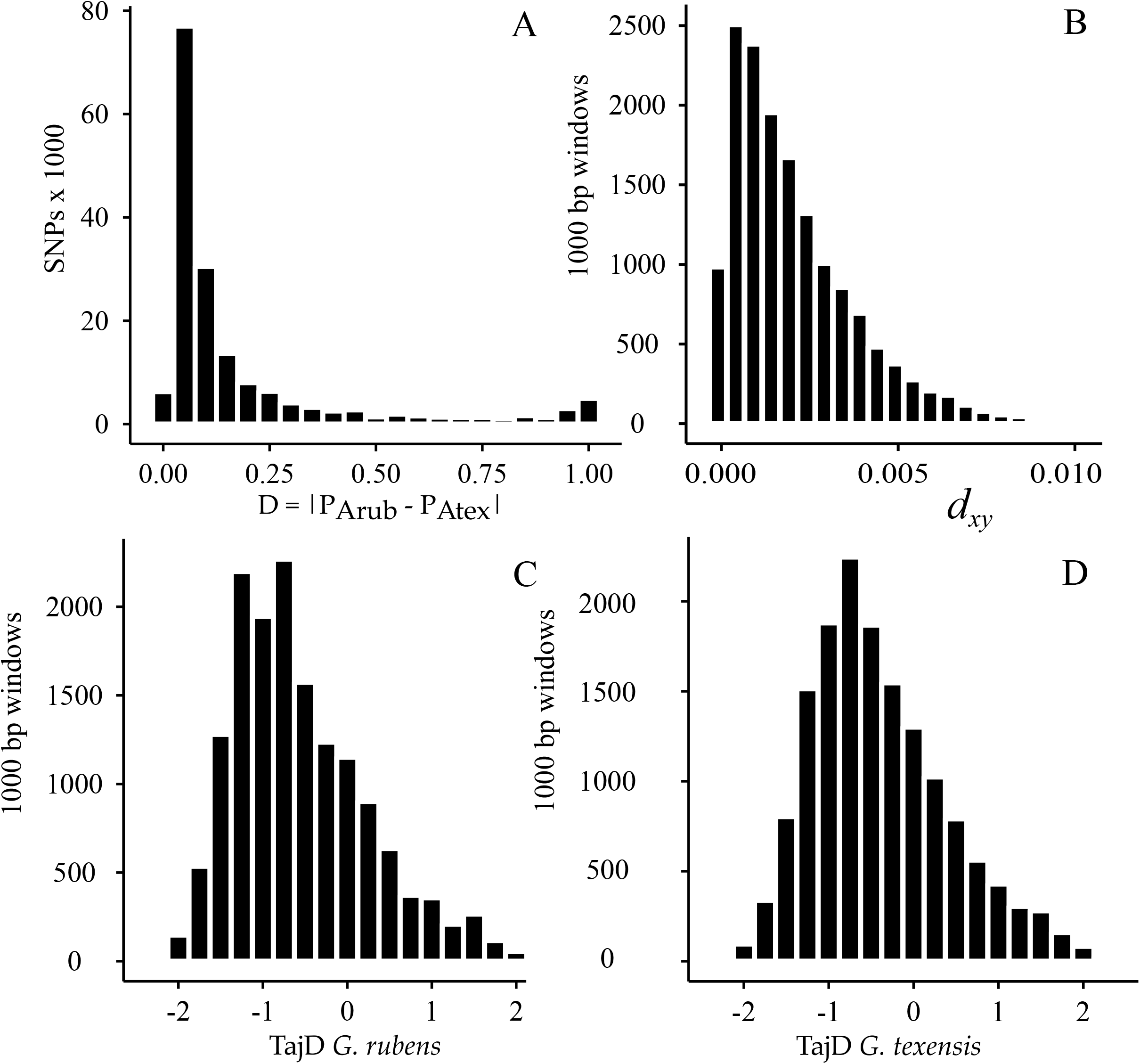
Genomic divergence. The distribution of the interspecific allele frequency difference, D, across SNPs (A), of the absolute divergence, *d_xy_*, in 1000 bp windows (B), and of Tajima’s D in 1000 bp windows for *G. rubens* (C) and *G. texensis* (D), respectively

### The demographic history

We found no substantial evidence for genetic structure in the two populations considered within either species. The species axis was the predominant axis of variation among individuals in the Principal Component Analysis (23.93% of total SNP variation, Fig. S1A), followed by axes separating *G. texensis* (PC2, 6.13% and PC3, 4.60%) and *G. rubens* (PC4, 4.35%) individuals. Variation within species was not related to geographic locations from which the individuals were collected (Fig. S1B, C). STRUCTURE further supported the finding that neither of the species was strongly differentiated geographically. The optimal K equaled 2 when we ran STRUCTURE with both species included (Fig. S2). Examining population structure within species revealed weak evidence for population substructure in both species at K=2, but K = 1 was the most parsimonious given the spread in log-likelihoods across K-values (Fig. S2). These results are robust across different subsets of SNPs and sample sizes (Fig. S3).

To infer the role of gene flow and bottlenecks during the evolutionary history of *G. texensis* and *G. rubens*, we used a nested rejection procedure to select the best model out of eight different models varying in the presence and timing of bottlenecks and gene flow (Fig. 3). First, we compared the gene flow models with each other. The gene flow model with the highest posterior probability was the ‘ancestral gene flow’ model (AGF P_posterior_ = 0.99 versus continuous gene flow, CGF: P_posterior_ < 0.01, and recent gene flow, RGF: P_posterior_ =0.01). Then we compared the bottleneck models with each other and found that the *‘G. rubens* bottleneck’ model had the highest posterior probability (RB P_posterior_ = 0.67 versus *G. texensis* bottleneck, TB: P_posterior_ = 0.43 and both bottleneck, BB: P_posterior_ < 0.01). We then combined these best models into a model with both ancestral gene flow and a bottleneck for *G. rubens* (AGFRB) and compared that model against a simple divergence model (DIV), the best gene flow model (AGF), and the best bottleneck model (RB). In this final model comparison, the combined model had the highest posterior probability (AGFRB: P_posterior_ = 0.68; AGF: P_posterior_ = 0.22; DIV: P _posterior_ = 0.02; RB: P_posterior_ = 0.08; Fig. 3, Fig. 4). Similar results were obtained using the full sample, including additional, but potentially related individuals: AGFRB: P_posterior_ = 0.75; AGF: P_posterior_ = 0.21; DIV: P_posterior_ = 0.03; RB: P_posterior_ = 0.01.

**Fig. 3.**
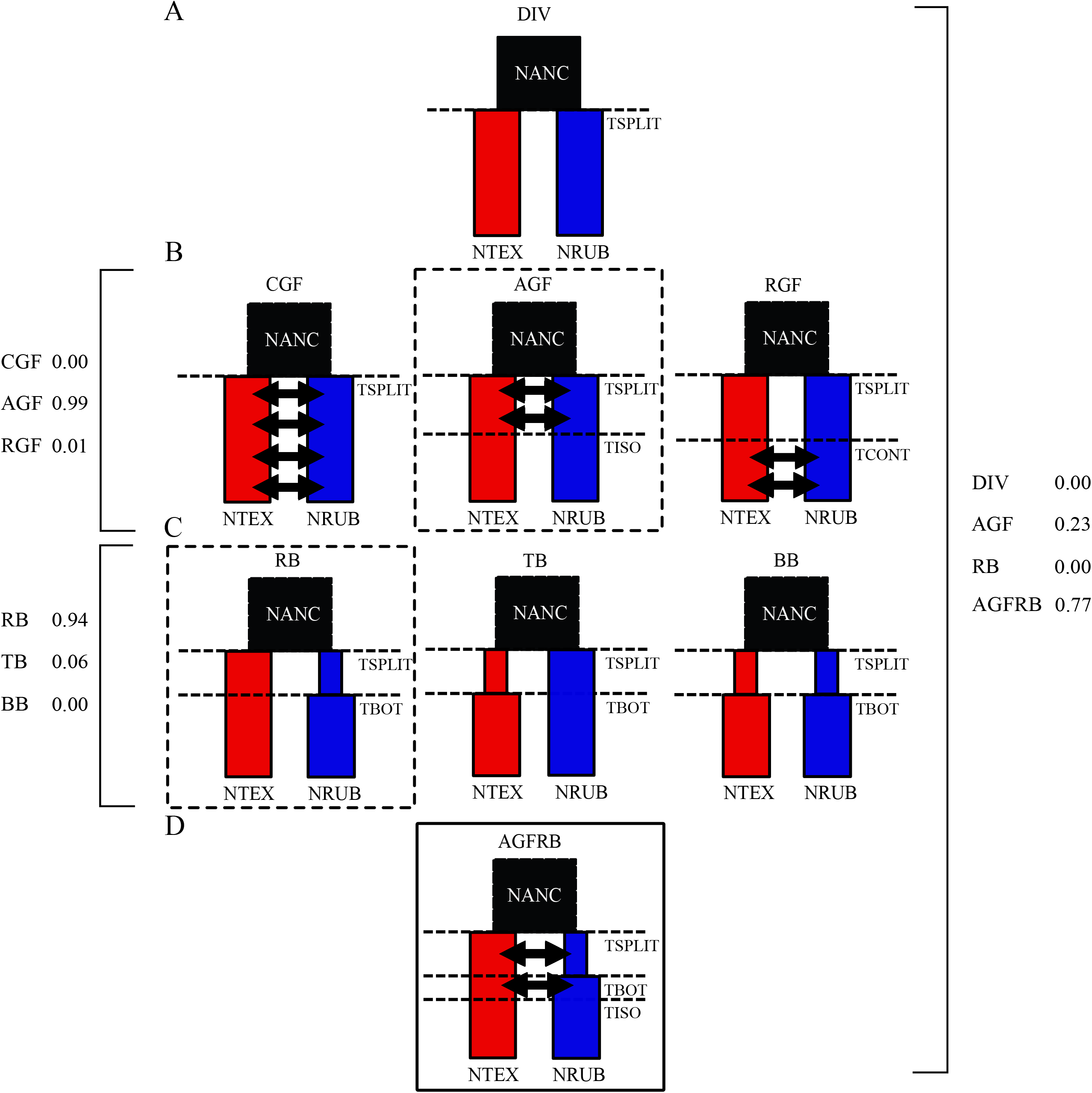
Demographic scenarios for Approximate Bayesian Computation. Eight scenarios were simulated under the ABC framework. (A) A simple divergence scenario (DIV) with a log uniform prior on the divergence time (T_SPLIT_), the ancestral population size (N_ANC_) and the current effective population sizes for *G. rubens* and *G. texensis* (N_RUB_, N_TEX_). (B) Three different gene flow models with either continuous gene flow (CGF), ancestral gene flow (AGF), or recent gene flow (secondary contact; RGF) were additionally defined by parameters describing migration rates (M_TEX>>RUB_, M_RUB>>TEX_; uniform priors not overlapping zero) and the time point since cessation of gene flow (T_ISO_) or of secondary contact (T_CONT_), both with log uniform priors. (C) Three bottleneck models defined by the time since recovery to current population sizes (TBOT; log uniform prior) and the relative population size reduction (BOTSIZE; uniform prior not overlapping zero) for *G. rubens* (RB), *G. texensis* (TB), or both (BB). (D) An additional model (AGFRB) combining the best gene flow (AGF) and best bottleneck (RB) model, marked by the black, dashed rectangles. The posterior probabilities for model selection are given left of the square (opening) brackets for the three gene flow and the three bottleneck models, and right of the square (closing) brackets for the final model selection step.

**Fig. 4.**
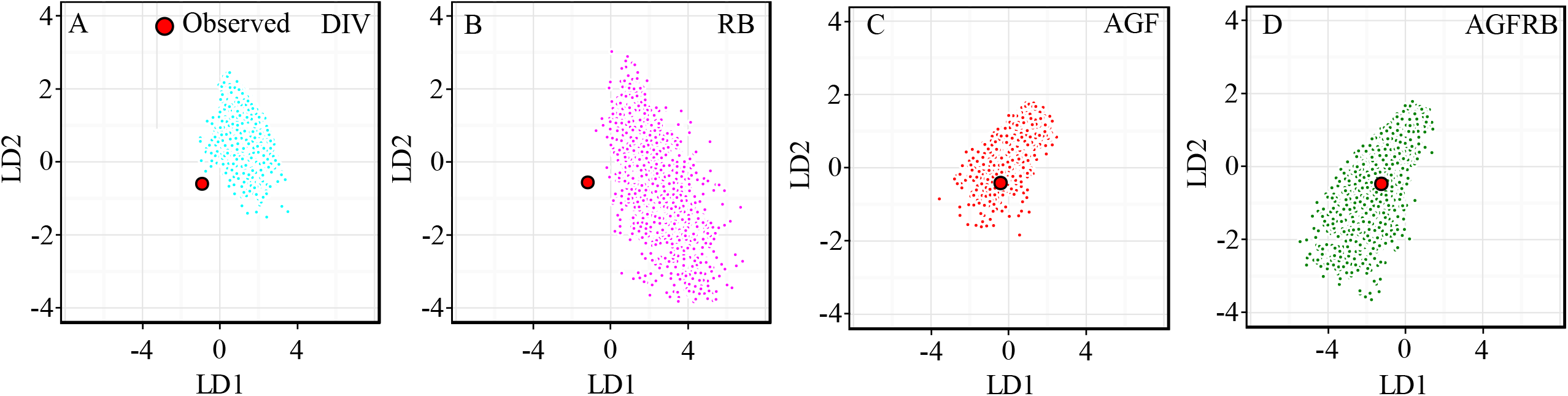
Distribution of observed and simulated data sets in multivariate summary statistic space. For each of the four models used in the final model selection step (see also Fig. 3) the distribution of the 1% posterior samples with the smallest Euclidean distance to the observed data is shown relative to the coordinates of the observed data. The multivariate summary statistic space is constrained by the first two linear discriminants representing linear combinations of the summary statistics used in model selection (see text for details).

As posterior probabilities may differ even among very similar models, it is critical to evaluate statistical support for model choice. Overall, model choice was well supported. For each selection step, we used cross validation to verify that models can be distinguished by assuming one of the models is the ‘true’ model and then performing 1,000 independent model selection steps under that assumption. The accuracy with which the assumed ‘true’ model was chosen was high for the gene flow models (98%, 96%, and 52% for AGF, CGF, and RGF, respectively), bottleneck models (76%, 66%, and 71% of the time for RB, TB, and BB respectively), and the final model selection step (75%, 82%, 82%, 84% for DIV, AGF, RB, AGFRB, respectively). It is important to note that the AGFRB model had the highest support overall and final model selection was well supported, but there is overlap of the posterior distribution of the summary statistics in multivariate space between the AGF and AGFRB models (Fig. 4).

Because there was some overlap between the posteriors of AGF and AGFRB (Fig. 4), and AGFRB only differs from AGF in the addition of a bottleneck, both models were used for demographic parameter estimates. Divergence times were distributed rather widely in both the AGF and AGFRB scenario and posterior density distributions were widely overlapping., The median divergence time varied between 350,000 years ago (700,000 generations ago) for AGF and double that for AGFRB. The ancestral effective population size was estimated around 200,000, almost an order of magnitude higher than the model estimates for current effective population sizes in *G. rubens* (~31,000 for AGFRB and ~18,000 for AGF) and *G. texensis* (~60,000 and ~28,000; Table 1, Table S2, Fig. 6A). A bottleneck for *G. rubens* was estimated at 15% of the current effective population size (Table 1, Fig. 6C) and recovery to current population sizes was achieved around 50,000 years ago (Table 1, Fig. 6B). Ancestral gene flow was bidirectional (median *m* = 0.18 and *m* = 0.27 for gene flow from *G. texensis* into *G. rubens* and vice versa, respectively; Table 1, Fig. 6C) and ceased around 18,000 years ago (Table 1, Table S2, Fig. 6B). The parameter estimates for the main model, AGFRB, were robust to the inclusion of additional, but potentially related, individuals; the estimates for times and population sizes were slightly higher and the inclusion of more samples gave similar results but at slightly higher accuracy (narrower HPD interval, Table S3, Fig. S4).

Statistical support for parameter inference varied across demographic events. Overall, the observed summary statistics fell well within the range of the simulated multivariate summary statistics under the AGF and AGFRB models (Fig. 4) and 95% HPD intervals of the distributions were generally narrow (Fig. 6, Table 1). For some demographic parameters (current population sizes for *G. rubens* [N_RUB_] and *G. texensis* [N_TEX_], and time since cessation of gene flow [T_ISO_] support was high (R^2^ > 0.81; RMSEP < 0.44); for other parameters estimated error rates were appreciably higher (Table 1, Table S2).

We compared *F_ST_* distributions simulated under the AGF, CGF, RGF, and AGFRB models with the observed *F_ST_* distribution as a measure of the effect of demography on the patterns of transcriptome-wide genetic variation. We found that the observed distribution (red line in Fig. 5) closely matched the simulated distribution of the two models with ancestral gene flow for most parts, including the secondary peak at the highest *F_ST_* bin (0.95 < *F_ST_* ≤ 1.00, Fig. 5C, D). In contrast, the observed *F_ST_* distribution showed substantial mismatch with the recent and continuous gene flow models.

**Fig. 5.**
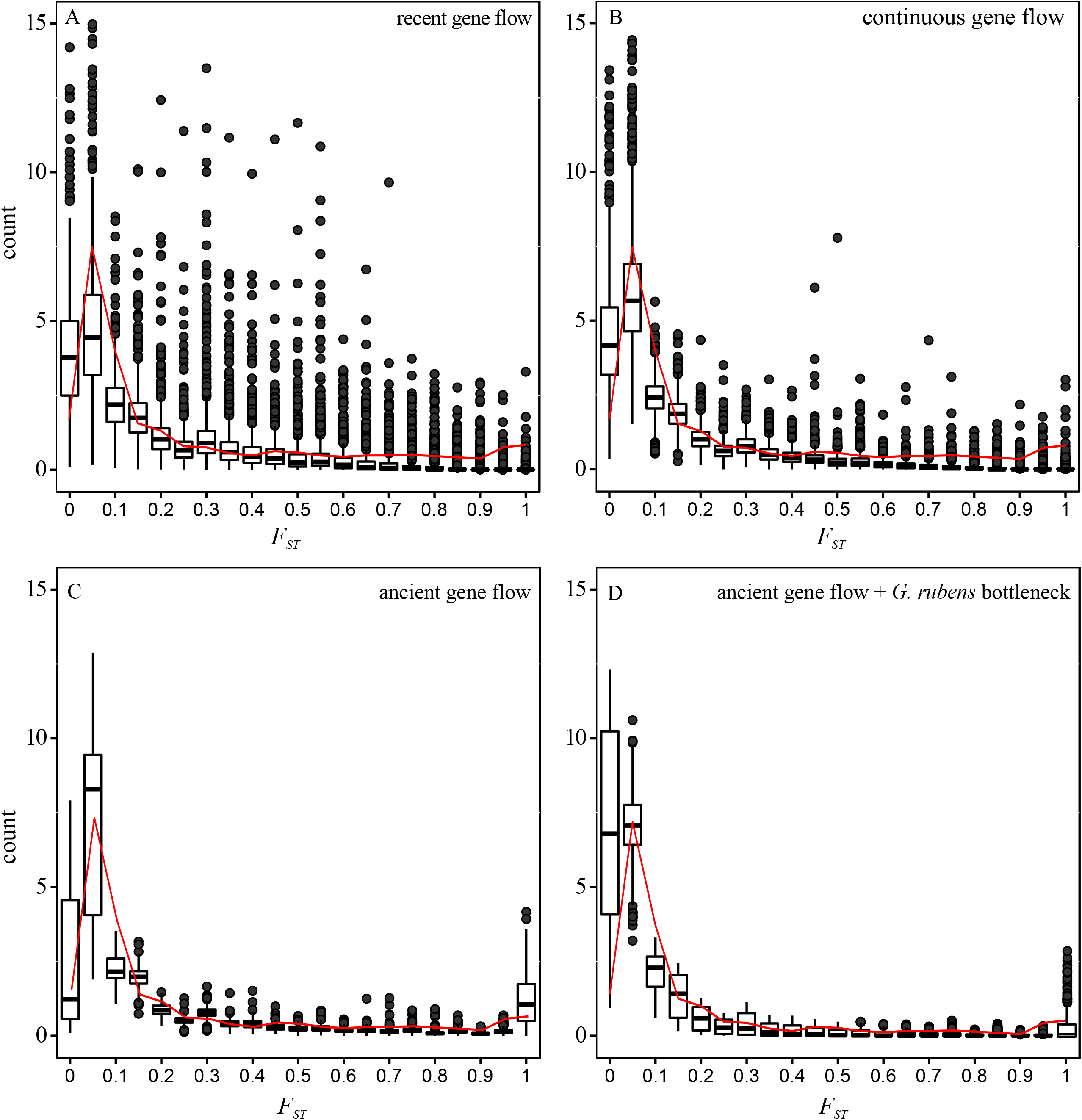
*F_ST_* distributions of simulated and observed data. The distribution of Weir and Cockerham’s *F_ST_* as calculated by the program arlsumstat are shown for 2,000 simulated data sets for different demographic model: recent gene flow (secondary contact; RGF), continuous gene flow (CGF), ancestral gene flow (AGF), and the AGFRB model. Observed data (1,000 1kbp sequences) are represented by the red solid line. The histograms show the density (y-axis) to enhance comparison between simulated and observed data.

### The role of selection

The *F_ST_* approach gave by far the highest number of outlier contigs. There were 514 contigs (5.8% of contigs) that had at least one SNP designated as a selection outlier (99^th^ quantile) in Arlequin’s *F_ST_* based hierarchical island method. There were no significantly (FDR < 10%) enriched Gene Ontology categories among the predicted gene products of these contigs and the most strongly enriched categories included mitochondrial processes, GTPase activity and cellular metabolism (Table S4, Table S5, Fig S5).

There were 80 contigs with *d_xy_* values in the 99^th^ percentile. The putative gene products corresponding to these 80 contigs were significantly (FDR < 10%) enriched for pheromone biosynthesis, hormone biosynthesis, mating behavior, and protein maturation (Table S4). Several of the most divergent loci match genes involved in *Drosophila melanogaster* sex pheromone pathways, such as *α-esterase* and *Desaturasel*, mushroom body development and neuromuscular synaptic targets, such as *S-lap1, tartan*, including those involved in flight muscle activity *(Stretchin-Mlck)*, and acoustic mating behavior, such as *Juvenile hormone esterase* and *calmodulin* (Table S6).

We retained 55 and 92 contigs that showed possible signatures of recent selective sweeps (Tajima’s D below 5% of the simulated sequences under the AGFRB scenario and π and *F_ST_* in the 90^th^ percentile) in *G. texensis* and *G. rubens*, respectively. The combined set of outlier loci was not significantly enriched for any biological processes after FDR correction. The most strongly enriched GO terms were predominantly higher order GO terms such as ‘organelle organization’, ‘primary metabolic process’, and ‘regulation of biological process’, but also contained more specific terms: ‘sperm mitochondrion organization’, ‘oocyte fate determination’, and ‘regulation of female receptivity’ (Table S4). Six contigs were shared between the species-specific sets of loci that showed potential signatures of a recent selective sweep signature. Three of these have no functionally characterized gene products. The other three are *neuroglian* (nrg), which is involved in various aspects of nervous system development and associated with male and female courtship behavior in *D. melanogaster; discs large 1* (dlg1), which affects neuromuscular junctions and changes fruit fly behavior across several domains including circadian activity and courtship; and *secretory 23* (sec23), which is an important component in differentiation of extra-cellular membranes in neurons and epithelial cells (Table S7). Several other gene products associated with contigs in the species-specific sets have functional roles in calcium or potassium channel activity (*e.g*., *nervana2*, expressed in the *Drosophila* auditory organs), nervous system development (*e.g*. *muscleblind*, which also alters female receptivity during courtship), veined-wing song generation (*e.g*. *period)*, as well as many genes related to metabolic and cellular processes.

There was one (unannotated) contig shared between the *d_xy_* approach and the selective sweep approach. Additionally, among the 514 outlier loci detected in Arlequin encompassed 11 contigs also found with the *d_xy_* approach, and 25 and 9 contigs respectively that were shared with the *G. rubens* and *G. texensis* specific selective sweep approach. These included the genes described above that are potentially related to sex pheromones biosynthesis (Desat1), flight muscle activity (Mlc-k), sensory neuron development (nrg), and auditory pathway ion channel activity (nrv2).

## DISCUSSION

Here, we illuminate the role of demographic and selective processes in shaping genetic variation during speciation. Combined insight in putative neutral (neutral divergence given the demographic history) and selective effects allowed us to infer the evolutionary history of *Gryllus rubens* and *G. texensis*, sibling species with large, overlapping distributions and strong phenotypic divergence in sexual traits with limited divergence in other phenotypes. We find strong support for a long history of ancestral gene flow and a bottleneck in *G. rubens*. Importantly, our data lend support to the hypothesis that loci showing high relative genetic differentiation compared to the genomic background may have evolved in response to demographic events and drift rather than in response to election. Interestingly, several of the loci with show signatures of positive or divergent selection after taking into account the effects from demography are potential orthologs of *D. melanogaster* genes involved in premating isolation, a major source of reproductive isolation between *G. rubens* and *G. texensis*. This work represents an important first step in assessing the contribution of neutral and selective forces to genetic divergence in a model system for sexual selection research.

### Neutral divergence and demography

We sequenced the transcriptomes of 40 individuals across four populations. Our observed transition:transversion ratio of 1.6:1 compares well with the estimate (1.55) from another cricket species pair, *G. firmus* and *G. pennsylvanicus* (Andrés et al. 2013), and suggests that sequencing errors did not contribute unduly to SNP discovery. Divergence across ~175K SNPs showed a bimodal and slightly right-skewed distribution of absolute (allele frequency) divergence, *D* (Fig. 2), and genetic differentiation, *F_ST_* (Fig. 5). The *F_ST_* distributions simulated under our top two scenarios were also right-skewed and strongly resembled the observed distribution of genetic differentiation, in strong contrast to *F_ST_* distributions corresponding to other models. Most importantly the simulated distributions under the most likely demographic scenarios, AGF and AGFRB, showed secondary peaks at *F_ST_* > 0.95. This indicates that a significant proportion of our fixed loci may have risen to fixation stochastically due to neutral processes (a combination of drift, population size variation, and gene flow) while gene flow homogenizes other (random) parts of the genome. Concordantly, the *F_ST_* based approach uncovered substantially more loci with putative signatures of positive selection than methods based on allele frequency spectra (with thresholds informed by inferred demographic history) or absolute sequence divergence (514 contigs in the *F_ST_* approach versus ~50-90 contigs in the other approaches). Our findings emphasizes the shortcomings of traditional *F_ST_* outlier approaches to discern selection effects from genomic background variation (Narum and Hess 2011; Lotterhos and Whitlock 2014).

We find strong evidence for a long history of bidirectional gene flow before *G. rubens* and *G. texensis* became fully reproductively isolated around 18,000 years ago, sometime during the last Pleistocene glacial cycles. This finding adds to a growing body of work that suggest divergence can occur in the face of gene flow (Bolnick and Fitzpatrick 2007; Nosil 2008; Bird et al. 2012; Feder et al. 2013). A large amount of recent work has focused on the role of gene flow in speciation, especially in combination with divergent or positive selection. In the genic view of speciation (Wu 2001) most areas of the genome are homogenized among populations during divergence with gene flow, and regions showing excess differentiation are thus likely protected by selection. This idea has been tested in many model systems with mixed results (Turner et al. 2005; Ellegren et al. 2012; Nosil et al. 2012; Cruickshank and Hahn 2014; Burri et al. 2015; Marques et al. 2016). Recent work suggests that genomic mosaics may in fact be mostly a consequence of linked selection caused by differences in recombination rates and density of selected loci and are thus expected to be conserved in pairwise comparisons even among distantly related taxa (Nachman and Payseur 2012; Burri et al. 2015; Van Doren et al. 2017). Our results support this idea as our demographic simulations recreated heterogeneous patterns similar to our observed data. Although selection certainly contributed to transcriptome divergence in *G. rubens* and *G. texensis* our results suggest a larger role for neutral divergence shaped by the effects of migration and population size variation and echo recent insights into the importance of considering neutral divergence when interpreting potential selection effects (*e.g*. reviewed in Ravinet et al. 2017).

In addition to bi-directional gene flow, the early stages of divergence between *G. texensis* and *G. rubens* were also influenced by a substantial bottleneck in *G. rubens*. There is some overlap between the AGF (no bottleneck) and AGFRB (with a *G. rubens* bottleneck) scenarios in the simulated summary statistic distribution, but the latter has a substantially higher posterior probability and corroborates the peripatric origin for *G. rubens* hypothesized in a previous study (Gray et al. 2008). Although that study used a single mitochondrial locus, it was done with extensive geographic sampling, and both studies suggest a bottleneck for *G. rubens*. Furthermore, estimates of strong admixture between populations within species and divergence time estimates are overlapping (this study: median ~ 0.35 - 0.70 million years ago; Gray *et al*. study: 0.25 – 2.0 mya). Estimates for current effective population sizes (roughly between 30 and 60 thousand for the AGFRB model and between 20 and 30 thousand for the AGF model) are surprisingly low given the potential census population size for *G. texensis* is in the millions (Gray et al. 2008). Potentially, the discrepancy is due to recent population expansion (Ptak and Przeworski 2002; Nadachowska-brzyska et al. 2013) or variation in individual mating success (Lande and Barrowclough 1987), as is observed in wild populations of closely related species (Ritz and Köhler 2010; Rodriguez-Munoz et al. 2010).

### τhe role of selection

A central aim of this study was to elucidate the role of selection during divergence within the context of the inferred demographic history. The species have strongly divergent mating behaviors with no evidence for reinforcement (Gray and Cade 2000; Higgins and Waugaman 2004; Izzo and Gray 2004; Blankers et al. 2015a). Many other cricket species show similarly strong divergence in various aspects of their mating behavior and several lines of evidence from various taxa indicate that this is at least in part driven by selection (Gray and Cade 2000; Bentsen et al. 2006; Shaw et al. 2007; Bailey 2008; Thomas and Simmons 2009; Oh and Shaw 2013; Blankers et al. 2017; Pascoal et al. 2017). Here, we show that the striking behavioral divergence is to some extent reflected in elevated sequence divergence of loci with putative functions in acoustic and chemical mating behavior. We find evidence that the set of loci showing the highest levels of sequence divergence are enriched for contigs bearing significant similarity to genes with known function in mating behavior in *D. melanogaster*. In addition, among the six contigs that showed evidence for a selective sweep in both species, three are potential orthologs of genes that affect neuromuscular properties in fruit flies and have effects on the flies’ mating behavior. Several other species-specific outliers are potential orthologs of genes that can be tied to mating behavior variation in *Drosophila spp*.

Given the substantial time since divergence and the long history of gene flow, high sequence divergence is expected for loci that have experienced limited homogenizing effects from gene flow relative to the rest of the genome. The theoretical support for speciation with gene flow driven by divergence in secondary sexual characters is very thin at best (van Doorn et al. 2004; Weissing et al. 2011; Servedio 2015). Here we provide exciting and rare evidence for speciation with primary gene flow while both phenotypic (Gray and Cade 2000), quantitative genetic (Blankers et al. 2015b, 2017), and genomic analyses (this study) highlight a role for selection on (acoustic) mating behavior in driving reproductive isolation. A compelling alternative interpretation of the findings here is that the peripatric origin of *G. rubens* has allowed for an initial phase of reduced gene flow; during this phase mating signals and preferences may have diverged sufficiently (aided by a founder effect following a population bottleneck) to maintain reproductive isolation during a subsequent phase of range expansion culminating into the contemporary, widespread, and largely overlapping species’ distributions. More empirical studies examining the role of gene flow and selection in systems characterized by strong sexual isolation are needed to test the theoretical predictions for speciation by sexual selection. However, this study along with other recent findings in finches (Campagna et al. 2017), fresh water stickleback (Marques et al. 2017), and cichlids (Malinksy et al. 2015) provide exciting first genomic insights into the joint effects from mating behavior divergence, sexual selection, and gene flow in the earliest phases of speciation.

We acknowledge that there are likely to be false positives among the detected outliers, as both linked (background) selection and demographic effects are expected to confound the signatures of positive or divergent selection (Cruickshank and Hahn 2014; Ravinet et al. 2017) and *a priori* expectations also increase the risk of “storytelling” (Pavlidis et al. 2012). By using coalescent simulations under the inferred evolutionary history, we have accounted for some confounding effects from demography. However, there is still potential neutral genetic variation that is unaccounted for, most notably the potentially confounding effects of recent population expansion and variation in recombination rates. We therefore caution that there is the uncertainty associated with the results obtained here and with genomic scans on quantitative traits in general (Jiggins and Martin 2017). Nevertheless, our findings provide exciting incentive for validation using alternative methods (*e.g*., QTL mapping) and follow-up functional genomic analyses.

Unsurprisingly, not all “outlier” contigs could be linked to mating behavior. The rest of these outliers are likely comprised of three groups: (1) Loci that are physically linked to loci under selection: In the earliest phases of speciation, only loci directly under strong divergent selection will differ. However, gene frequencies at tightly linked loci will also change and, given sufficient time as well as low to moderate migration and recombination rates, these loci will be swept to fixation along with selected sites (Smith and Haigh 1974) in a process called divergence hitchhiking (Feder et al. 2012; Via 2012); (2) Loci that are under selective forces that we have not yet elucidated: It is unlikely that divergent selection only targets loci involved in mating behavior and other traits may be differentiated between *G. rubens* and *G. texensis*. For example, females differ in the length of the ovipositor (Gray et al. 2001), a trait which reflects potential ecological adaptation to different soil types (Bradford et al. 1993); (3) Loci that are not under selection: Genetic drift can cause loci to drift to fixation and demographic effects such as bottlenecks and migration patterns (Holsinger and Weir 2009) can aid this process. Our simulations predict a significant number of fixed loci (1.90% on average for the AGFRB scenario) solely due to neutral processes (Fig. 5). Additionally, practical limitations of discovering low-frequency SNPs causing ascertainment bias (Clark et al. 2005) can contribute to misinterpretation of the patterns of genetic diversity (Vitti et al. 2013). A genomic map of *Gryllus* and further analyses would make strong headway into determining which of these categories the other potential outliers fall into.

Finally, there may be loci that are under selection but that were not detected by our scan because they simply were not being expressed. We sequenced samples from first generation laboratory offspring rather than animals directly from the field. Despite the fact that there are no differences between *G. texensis* and *G. rubens* in ecology, microhabitat use, or feeding behavior have been described (but note there is variation in the ovipositor length which is a potential adaptation to soil properties), the laboratory conditions have potentially limited our potential to detect genetic differences related to local adaptation.

In summary, this study underlines the importance of considering the joint effects from neutral divergence and selection in understanding the speciation process. Our results also offer unprecedented insight into the evolutionary history and the role of demography and selection in driving transcriptomic divergence in two sexually isolated field cricket sister species. We inferred that a long period of bidirectional, ancestral gene flow and a bottleneck in *G. rubens* preceded completion of reproductive isolation (Fig. 3,6). Importantly, the timing of gene flow appears to have significantly influenced the pattern of divergence (*i.e*. the *F_ST_* distribution) that we observe (Fig. 5). We also uncovered several loci that show signatures of positive or divergent selection and show that these contigs are potentially associated with courtship behavior, neuromuscular development, and chemical mating behavior. Future work will place these data on a genomic map allowing us to determine how genetic divergence is distributed relative to loci under selection. These findings provide important steps towards understanding the role of selective and neutral processes in shaping patterns of divergence and the role of sexual selection during speciation-with-gene flow. They also highlight the strength of combining information on (i) the phenotypes that contribute to reproductive isolation, (ii) demographic inference, and (iii) scans for loci under selection.

**Fig. 6.**
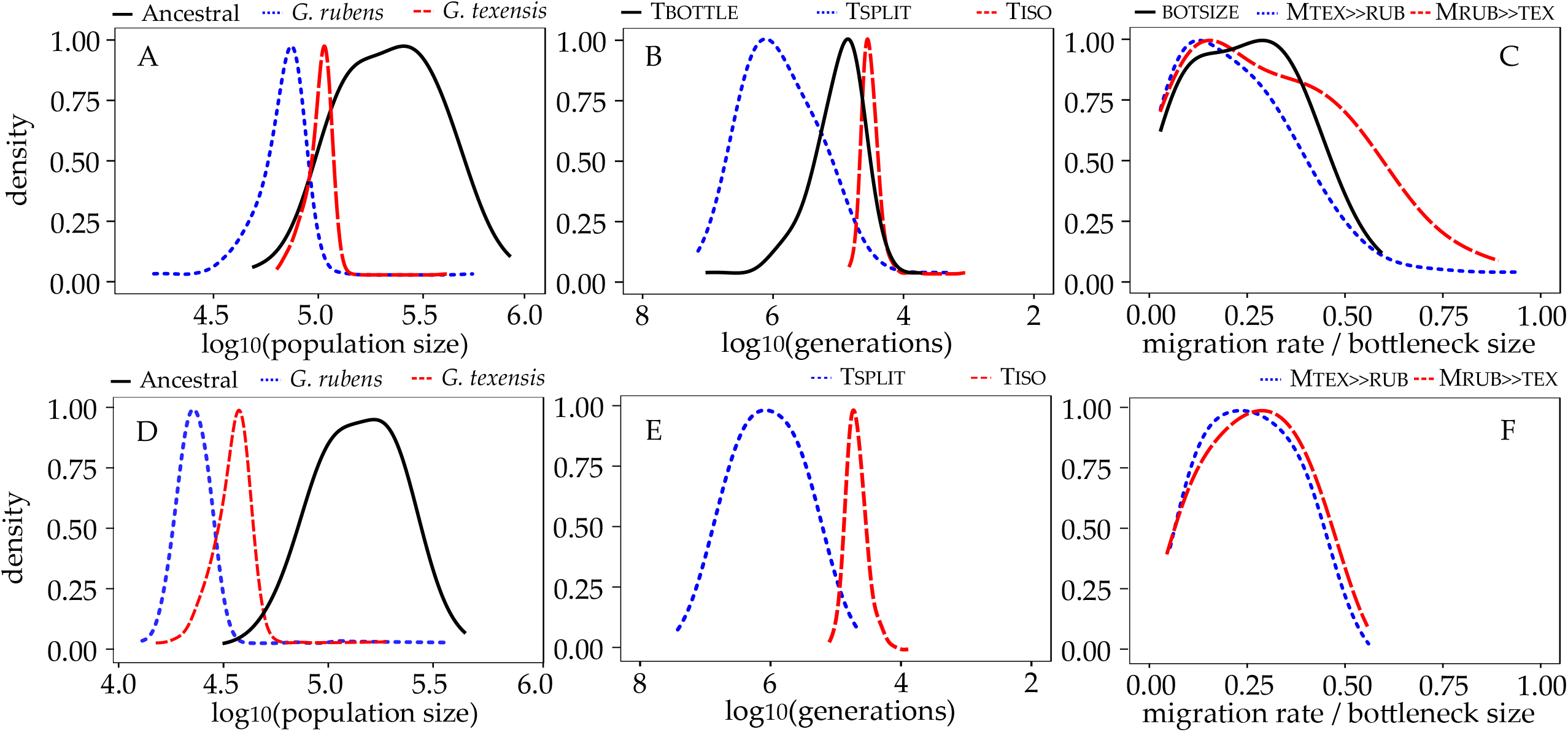
Demographic parameter estimation. For the AGFRB (A-C) and the AGF (D-F) models, the density distribution of the ancestral and current population sizes (A, D), the time since divergence, cessation of gene flow, and recovery to current population sizes after the bottleneck (B, E), and the migration rates and bottleneck size (C, F) are shown. The density lines have been trimmed to the existent parameter distribution (i.e., no density extrapolation) and have been smoothed by adjusting the bandwidth. For lines within one panel the same smoothing bandwidth has been used.

## CONFLICT OF INTEREST

The authors declare no conflict of interest, financial or otherwise.

## DATA ACCESSIBILITY

Data, including raw reads, sequences used for demographic analyses and SNP data files used in outlier analysis, will be made available on Dryad and the NCBI SRA archive prior to publication.

## SUPPLEMENTARY INFORMATION

Table S1. Individual RNA-seq read mapping statistics

Table S2. ABC estimates for the AGF scenario

Table S3. ABC estimates for the full sample (including 8 individuals from half-sib pairs), AGFRB scenario

Table S4. GO enrichment results for *F_ST_*, *d_xy_*, and selective sweep outliers.

Table S5. *F_ST_* outlier loci.

Table S6. *d_xy_* outlier loci

Table S7A,B Selective sweep outlier loci for *G. texensis* and *G. rubens*

Fig S1-S7. See figures for figure legends.

**Table S1.** Individual RNA-seq read mapping statistics. Mapping rates were calculated using bowtie2 with default parameters.

| Sample ID | Species | Population | Sex | Mapping rate |
| --- | --- | --- | --- | --- |
| 30037 rub | *G. rubens* | Ocala | f | 84.52% |
| 30038 rub | *G. rubens* | Ocala | f | 85.33% |
| 30039 rub | *G. rubens* | Ocala | f | 85.66% |
| 30040 rub | *G. rubens* | Ocala | f | 84.35% |
| 30041 rub | *G. rubens* | Ocala | f | 84.85% |
| 30057 rub | *G. rubens* | Lake City | f | 88.40% |
| 30059 rub | *G. rubens* | Lake City | f | 88.86% |
| 30060 rub | *G. rubens* | Lake City | f | 87.83% |
| 30061 rub | *G. rubens* | Lake City | f | 90.23% |
| 30052 rub | *G. rubens* | Ocala | m | 78.01% |
| 30053 rub | *G. rubens* | Ocala | m | 80.72% |
| 30055 rub | *G. rubens* | Ocala | m | 79.76% |
| 30063 rub | *G. rubens* | Lake City | m | 77.70% |
| 30064 rub | *G. rubens* | Lake City | m | 77.56% |
| 30065 rub | *G. rubens* | Lake City | m | 70.75% |
| 30027 tex | *G. texensis* | Lancaster | f | 83.09% |
| 30028 tex | *G. texensis* | Lancaster | f | 83.20% |
| 30029 tex | *G. texensis* | Lancaster | f | 81.61% |
| 30030 tex | *G. texensis* | Lancaster | f | 83.80% |
| 30031 tex | *G. texensis* | Lancaster | f | 80.42% |
| 30043 tex | *G. texensis* | Austin | f | 91.78% |
| 30044 tex | *G. texensis* | Austin | f | 90.01% |
| 30046 tex | *G. texensis* | Austin | f | 87.70% |
| 30032 tex | *G. texensis* | Lancaster | m | 76.17% |
| 30033 tex | *G. texensis* | Lancaster | m | 77.76% |
| 30034 tex | *G. texensis* | Lancaster | m | 77.24% |
| 30035 tex | *G. texensis* | Lancaster | m | 80.79% |
| 30036 tex | *G. texensis* | Lancaster | m | 76.77% |
| 30047 tex | *G. texensis* | Austin | m | 86.40% |
| 30049 tex | *G. texensis* | Austin | m | 88.52% |
| 30050 tex | *G. texensis* | Austin | m | 79.15% |
| 30051 tex | *G. texensis* | Austin | m | 86.18% |

**Table S2.** ABC estimates for the AGF scenario. Prior distributions (log-scale), posterior predictive checks and posterior parameter estimates (log scale, median and 95% highest posterior density interval) for the model are shown.

| Parameter | Prior<sup>a</sup> |  | Validation |  | Posterior |  |  |
| --- | --- | --- | --- | --- | --- | --- | --- |
|  | minimum | maximum | R<sup>2</sup> | RMSEP | 2.5% | Median | 97.5% |
| LOG<sub>10</sub>(N<sub>ANC</sub>) | 4.0 | 6.0 (lu) | 0.0 | 0.96 | 4.76 | 5.31 | 5.81 |
| LOG<sub>10</sub>(N<sub>RUB</sub>) | 3.0 | 6.0 (lu) | 0.93 | 0.27 | 3.81 | 4.26 | 4.55 |
| LOG<sub>10</sub>(N<sub>TEX</sub>) | 3.0 | 6.0 (lu) | 0.93 | 0.27 | 3.98 | 4.45 | 4.69 |
| LOG<sub>10</sub>(T<sub>SPLIT</sub>)<sup>b</sup> | 5.0 | 7.0 (lu) | 0.06 | 0.97 | 4.61 | 5.83 | 7.04 |
| LOG<sub>10</sub>(T<sub>ISO</sub>)<sup>b</sup> | 3.0 | 7.0 (lu) | 0.79 | 0.46 | 4.21 | 4.56 | 4.73 |
| M<sub>TEX>>RUB</sub> | 0.01 | 0.5 (u) | 0.17 | 0.91 | 0.03 | 0.24 | 0.49 |
| M<sub>RUB>>TEX</sub> | 0.01 | 0.5 (u) | 0.12 | 0.94 | 0.01 | 0.26 | 0.51 |

<sup>a</sup> priors are uniformly (u) or log-uniformly (lu) distributed and do not overlap zero for migration rates and bottleneck size.
<sup>b</sup> the timing of demographic events is in (logarithm of) number of generation and both species have two generations annually.

**Table S3.** ABC estimates for the full sample (including 8 individuals from half-sib pairs), AGFRB scenario. Prior distributions (log-scale), posterior predictive checks and posterior parameter estimates (log scale, median and 95% highest posterior density interval) for the model are shown.

| Parameter | Prior<sup>a</sup> |  | Validation |  | Posterior |  |  |
| --- | --- | --- | --- | --- | --- | --- | --- |
|  | minimum | maximum | R<sup>2</sup> | RMSEP | 2.5% | Median | 97.5% |
| LOG<sub>10</sub>(N<sub>ANC</sub>) | 4.0 | 6.0 (lu) | 0.05 | 0.974 | 4.94 | 5.32 | 5.72 |
| LOG<sub>10</sub>(N<sub>RUB</sub>) | 3.0 | 6.0 (lu) | 0.89 | 0.333 | 4.70 | 4.79 | 4.87 |
| LOG<sub>10</sub>(N<sub>TEX</sub>) | 3.0 | 6.0 (lu) | 0.88 | 0.346 | 4.73 | 4.85 | 4.94 |
| LOG<sub>10</sub>(T<sub>SPLIT</sub>)<sup>b</sup> | 5.0 | 7.0 (lu) | 0.01 | 0.997 | 5.49 | 6.23 | 6.74 |
| LOG<sub>10</sub>(T<sub>ISO</sub>)<sup>b</sup> | 3.0 | 7.0 (lu) | 0.81 | 0.438 | 4.27 | 4.53 | 4.72 |
| LOG<sub>10</sub>(T<sub>BOT</sub>)<sup>b</sup> | 5.0 | 7.0 (lu) | 0.02 | 0.990 | 5.14 | 5.19 | 5.32 |
| BOTSIZE | 0.01 | 0.5 (u) | 0.01 | 0.995 | 0.09 | 0.15 | 0.23 |
| M<sub>TEX>>RUB</sub> | 0.01 | 0.5 (u) | 0.12 | 0.938 | 0.05 | 0.12 | 0.18 |
| M<sub>RUB>>TEX</sub> | 0.01 | 0.5 (u) | 0.12 | 0.938 | 0.01 | 0.18 | 0.75 |

<sup>a</sup> priors are uniformly (u) or log-uniformly (lu) distributed and do not overlap zero for migration rates and bottleneck size.
<sup>b</sup> the timing of demographic events is in (logarithm of) number of generation and both species have two generations annually.

**Table S4.** GO enrichment results. The top ten terms of the Gene Ontology enrichment is shown for the *d_xy_* outliers and the Allele Frequency Spectrum (AFS) outliers. For each Biological Process, the number of annotated transcripts and the number of observed and expected transcripts in the sample with a given annotation are shown. The Fisher’s exact test P-value is corrected using the parent-child algorithm (Grossmann *et al*. 2007). The FDR is the false discovery rate based on the corrected P-values.

| GO | Term | #Annot | #Sample | #Exp | P-value | FDR |
| --- | --- | --- | --- | --- | --- | --- |
|  |  | ***Fst*** |  |  |  |  |
| GO:0032543 | mitochondrial translation | 10 | 4 | 0.34 | 0.00056 | 1 |
| GO:0043087 | regulation of GTPase activity | 54 | 10 | 1.85 | 0.00123 | 1 |
| GO:0071695 | anatomical structure maturation | 2 | 2 | 0.07 | 0.00141 | 1 |
| GO:0010822 | positive regulation of mitochondrion organization | 5 | 3 | 0.17 | 0.0026 | 1 |
| GO:0044267 | cellular protein metabolic process | 1773 | 74 | 60.58 | 0.00318 | 1 |
| GO:0022411 | cellular component disassembly | 56 | 7 | 1.91 | 0.00398 | 1 |
| GO:0000910 | cytokinesis | 248 | 19 | 8.47 | 0.00428 | 1 |
| GO:0043603 | cellular amide metabolic process | 577 | 30 | 19.72 | 0.00526 | 1 |
| GO:0030716 | oocyte fate determination | 58 | 6 | 1.98 | 0.00564 | 1 |
| GO:0050789 | regulation of biological process | 4028 | 160 | 137.63 | 0.00566 | 1 |
|  |  | ***d<sub>xy</sub>*** |  |  |  |  |
| GO:0042811 | pheromone biosynthetic process | 44 | 4 | 0.2 | 4.30E-06 | 0.0027 |
| GO:0042810 | pheromone metabolic process | 49 | 4 | 0.22 | 3.60E-05 | 0.0071 |
| GO:1903317 | regulation of protein maturation | 24 | 3 | 0.11 | 3.70E-05 | 0.0071 |
| GO:0042446 | hormone biosynthetic process | 82 | 4 | 0.37 | 4.50E-05 | 0.0071 |
| GO:1903318 | negative regulation of protein maturation | 23 | 3 | 0.1 | 0.0001 | 0.0152 |
| GO:0044705 | multi-organism reproductive behavior | 359 | 6 | 1.62 | 0.0002 | 0.0232 |
| GO:0019098 | reproductive behavior | 367 | 6 | 1.65 | 0.0005 | 0.0380 |
| GO:0007618 | mating | 400 | 6 | 1.8 | 0.0005 | 0.0380 |
| GO:0006551 | leucine metabolic process | 3 | 2 | 0.01 | 0.0011 | 0.0734 |
|  |  | ***AFS*** |  |  |  |  |
| GO:0006996 | organelle organization | 2271 | 41 | 21.7 | 0.0003 | 0.3545 |
| GO:1902589 | single-organism organelle organization | 1791 | 32 | 17.1 | 0.0004 | 0.3545 |
| GO:0044238 | primary metabolic process | 4836 | 59 | 46.2 | 0.0007 | 0.4181 |
| GO:0090066 | regulation of anatomical structure size | 375 | 12 | 3.6 | 0.0014 | 0.5867 |
| GO:0050789 | regulation of biological process | 4028 | 52 | 38.5 | 0.0025 | 0.5867 |
| GO:0030382 | sperm mitochondrion organization | 6 | 2 | 0.1 | 0.0027 | 0.5867 |
| GO:0065007 | biological regulation | 4463 | 56 | 42.7 | 0.0027 | 0.5867 |
| GO:0007294 | germarium-derived oocyte fate determination | 46 | 4 | 0.4 | 0.0028 | 0.5867 |
| GO:0030716 | oocyte fate determination | 58 | 4 | 0.6 | 0.0033 | 0.5867 |
| GO:0045924 | regulation of female receptivity | 7 | 2 | 0.1 | 0.0035 | 0.5867 |
| GO:0006996 | organelle organization | 2271 | 41 | 21.7 | 0.0003 | 0.3545 |

**Fig S1.**
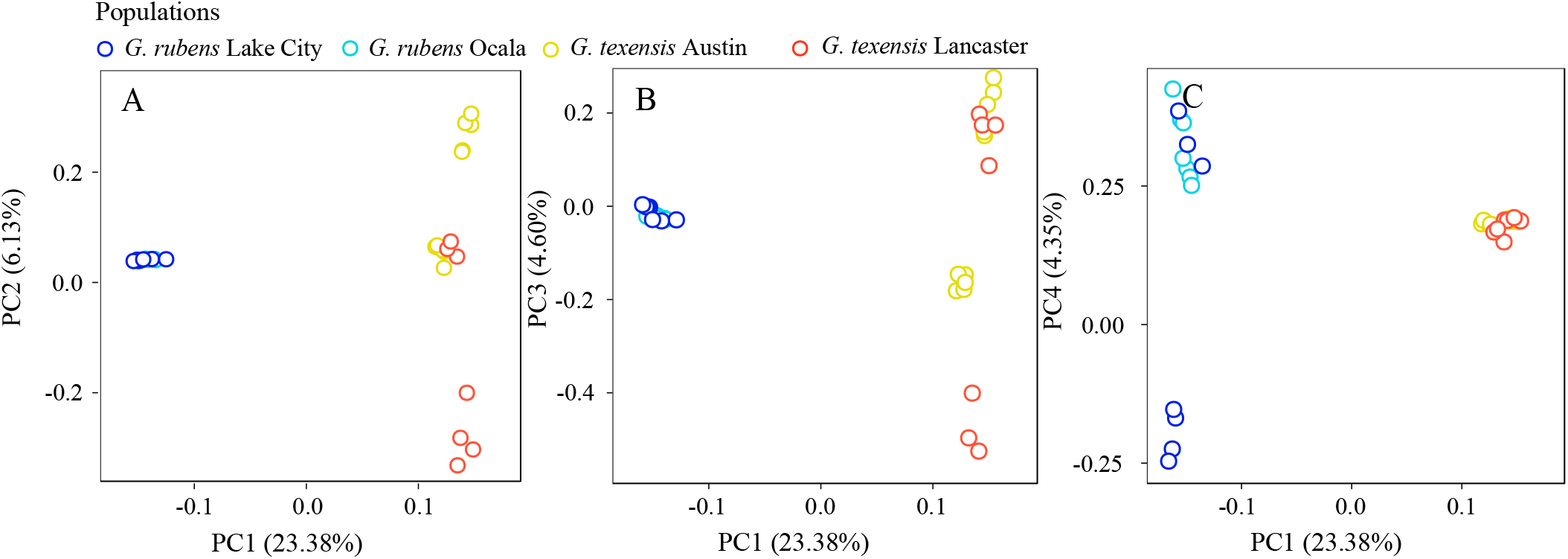
Population substructure in *G. rubens* and *G. texensis*. Variation in allele frequencies between species and between populations within species (Lake City and Ocala for *G. rubens*; Lancaster and Austin for *G. texensis)is* shown. The allele frequency variation in all 175,244 SNPs is summarized in the first four principal components teasing apart the species (PC1), and the populations in *G. texensis* (PC 2) and *G. rubens* (PC 4). Note that clustering along the PCs explaining within species variation among populations is much weaker compared to clustering of the species along PC1.

**Fig S2.**
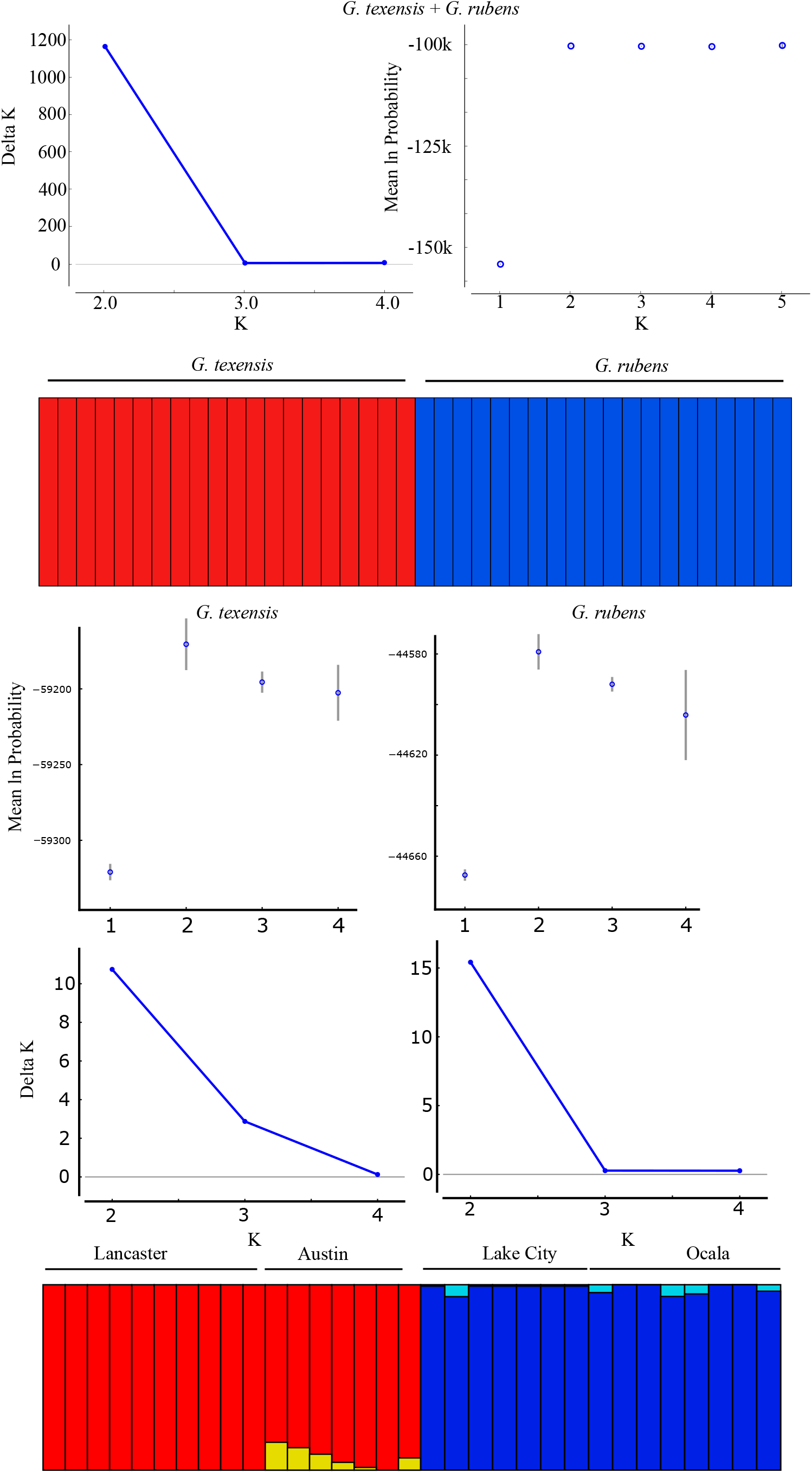
STRUCTURE results. For each of the species, STRUCTURE was run for 100,000 iterations at values for K=1 through K=4 (K=5 for the species combined). The mean natural logarithm of the probability and the delta K (increase or decrease in likelihood between consecutive runs for different values of K) were inspected to determine the most likely predicted number of populations. A run of *G. rubens* and *G. texensis* separately showed in both cases that, although the highest likelihood was for K=2 , differences with K=1 were only marginal and a defined pattern in population substructure was absent (see also the bar plots at the bottom). The run for the species combined (K=2) shows no introgression of *G. texensis* genes into the *G. rubens* or vice versa.

**Fig S3.**
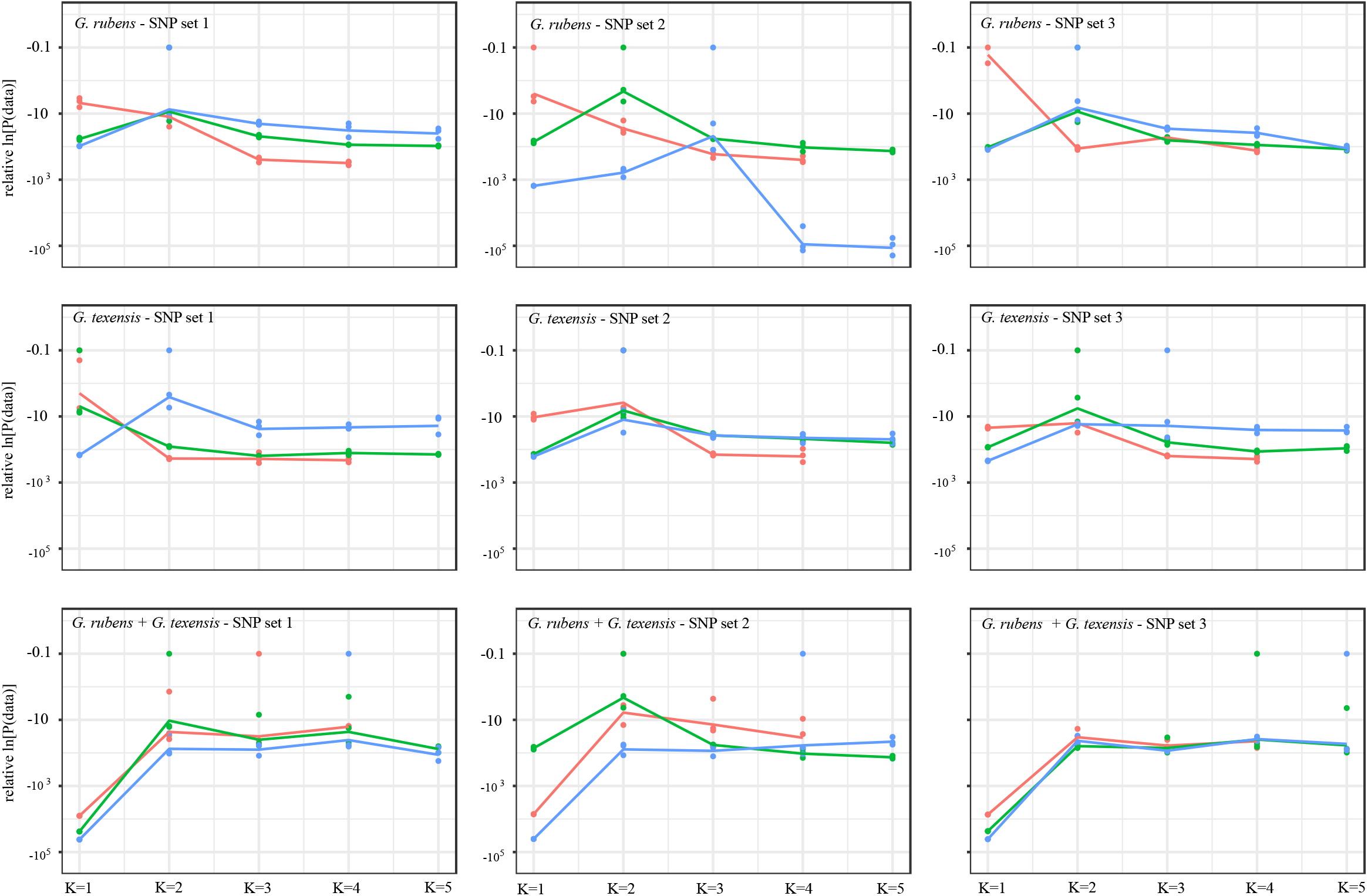
Relative natural log transformed probability of the data under different values for K. The raw probabilities from STRUCTURE relative to the maximum probability is shown for each K, for three random sets of 8835 SNPs (one per contig), and for *G. rubens, G. texensis*, and for the species combined (excluding eight individuals to correct for cryptic relatedness). Within each panel, the dots show each of the three iterations and the lines show the trend in the average difference in probability with the maximum probability for three different sample sizes: two random individuals per population (red), five random individuals per population (green), and all the individuals sampled from the populations.

**Fig S4.**
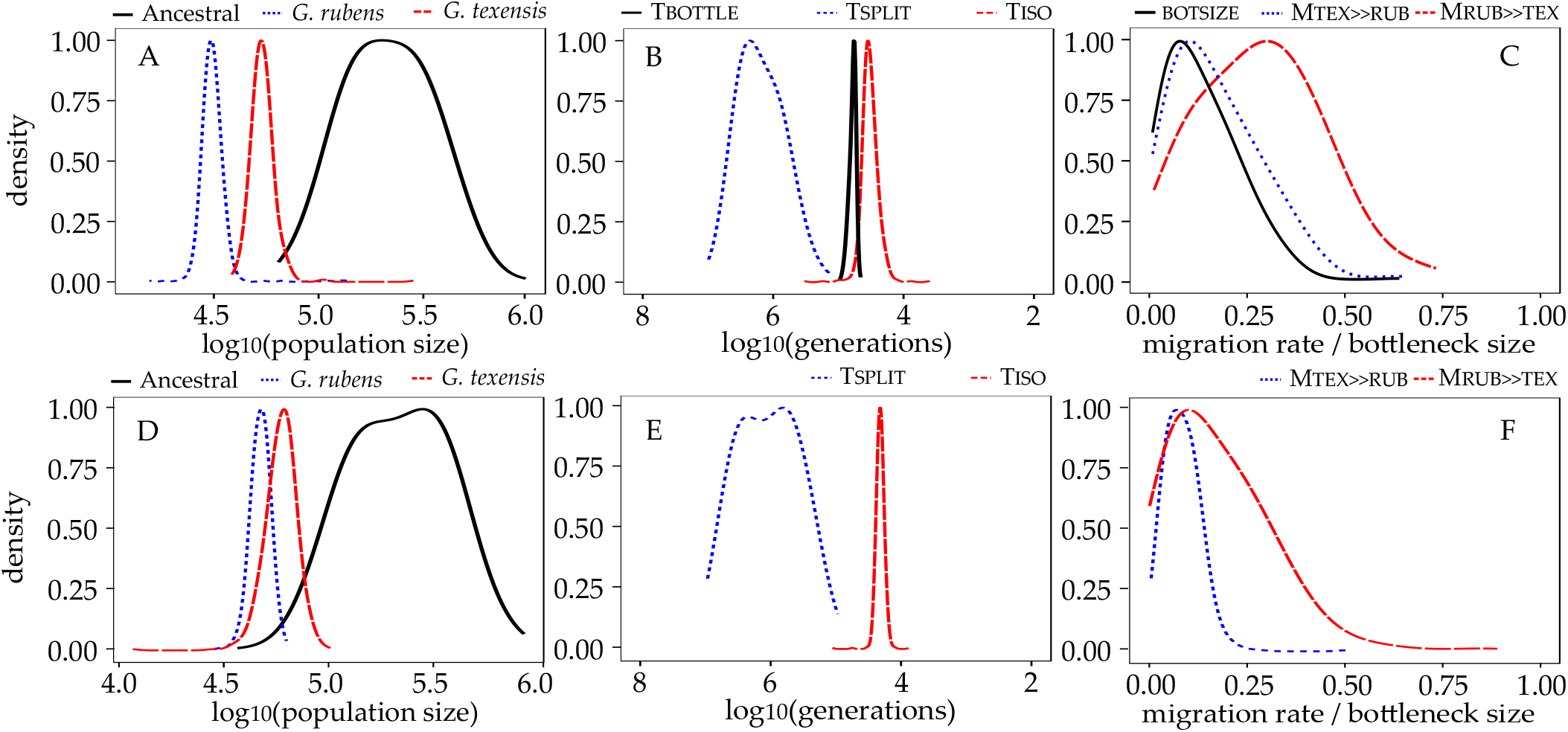
Demographic parameter estimation for the model with all 40 individuals. For the AGFRB (A-C) and the AGF models (D-F) the density distributions of the the ancestral and current population sizes (A,D), the time since divergence, cessation of gene flow, and recovery to current population sizes after the bottleneck (B,E), and the migration rates and bottleneck size (C,F) are shown. The density lines have been trimmed to the existent parameter distribution (i.e., no density extrapolation) and have been smoothed by adjusting the bandwidth. For lines within one panel the same smoothing bandwidth has been used.

**Fig S5.**
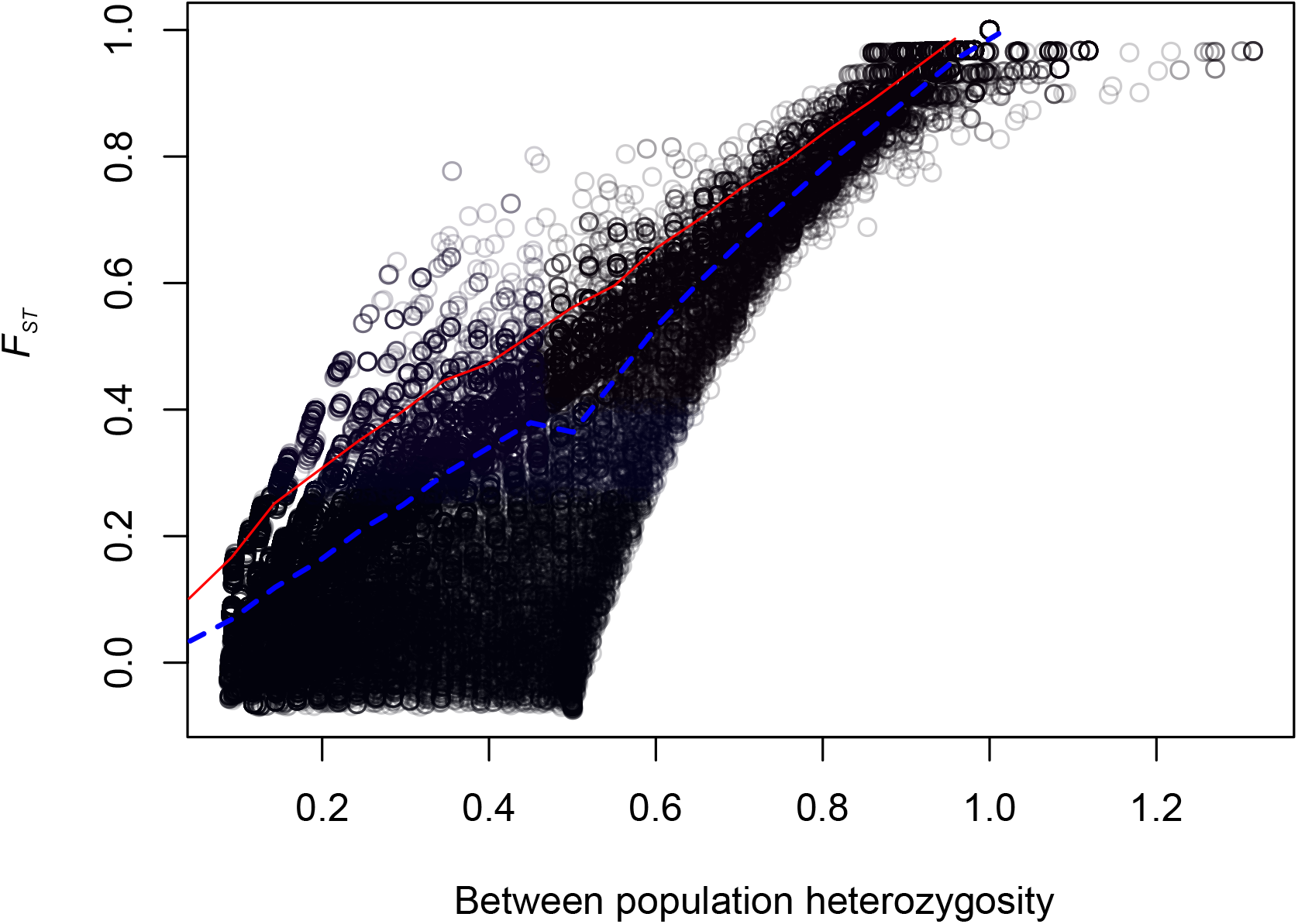
Arlequin FST based selection scan. The circles represent estimates for the FST and between population heterozygosity for all SNPs with MAF > 0.05 (81,125 SNPs). The blue dashed and red solid line are the median and 99th quantile, respectively, of the simulated null distribution for this relationship under a hierarchical island model. Any SNPs above the red solid lines were considered outliers.

## ACKNOWLEDGEMENTS

We thank Marie Jeschek and Harald Detering from the Berlin Center for Genomics and Biodiversity (BeGenDiv) for assistance in bioinformatics. We further thank Mike Ritchie, Roger Butlin, Daniel Wegmann, and Laurent Excoffier for helpful comments on the analyses and the Biostars and Stackoverflow community for help with scripting. The manuscript strongly benefited from the comments from the associate editor Jeffry Dudycha and three anonymous reviewers. Sample collection and processing comply with the “Principles of animal care”, publication No. 86-23, revised 1985 of the National Institute of Health, and also with the current laws of Germany. The authors declare no conflict of interest. This study is part of the GENART project. The work was supported by the Leibniz Association (SAW-2012-MfN-3). STV was supported by an Alexander von Humboldt Foundation fellowship.

